# *Leishmania* infection-derived extracellular vesicles drive transcription of genes involved in M2 polarization

**DOI:** 10.1101/2022.05.16.492154

**Authors:** Lisa Emerson, Anna Gioseffi, Hailey Barker, Austin Sheppe, Julianne K. Morrill, Mariola Edelmann, Peter Kima

**Affiliations:** Department of Microbiology and Cell Science, Institute of Food and Agricultural Sciences, University of Florida, Gainesville, Florida, United States of America

## Abstract

Although it is known that the composition of EVs is determined by the characteristics of the cell and its environment, the effects of intracellular infection on EV composition and functions are not well understood. We had previously shown that cultured macrophages infected with *Leishmania* parasites release EVs that contain parasite derived molecules (LiEVs). In this study we show that LdVash, a molecule previously identified in LiEVs from *L. donovani* infected RAW264.7 macrophages is widely distributed in the liver of *L. donovani* infected mice. This result shows for the first time that parasite molecules are released in EVs and distributed in infected tissues where they can be endocytosed by cells in the liver, including macrophages that undergo a significant increase in numbers as the infection progresses. To commence evaluating the potential impact of LiEVs on macrophage functions, we show that primary peritoneal exudate macrophages (PECs) express transcripts of signature molecules of M2 macrophages such as arginase 1, IL-10 and IL-4R when incubated with LiEVs. In comparative studies that illustrate how intracellular pathogens control the composition and functions of EVs released from macrophages, we show that EVs from RAW264.7 macrophages infected with Salmonella enterica serovar Typhimurium activate PECs to express transcripts of signature molecules of M1 macrophages such as iNOS, TNF alpha and IFN gamma and not M2 signature molecules. Finally, we show that in contrast to the polarized responses observed in in vitro studies of macrophages, both M1 and M2 signature molecules are detected in *L. donovani* infected livers although they exhibit differences in their spatial distribution in infected tissues.

## 1. Introduction

There is ample evidence that eukaryotic cells constitutively release extracellular vesicles (EVs) to their environment. EVs include exosomes that range in size from 30 -200 nm, microvesicles that are 50 -1000 nm, and apoptotic bodies from 500 -5000 nm [1]. In addition to the size differences between EVs, the vesicles also differ in their biogenesis and cargo content [1]. Although all EVs are likely to contribute to contactless cell-to-cell communication, numerous studies have focused on exosomes and their functions in these processes. There is increasing evidence that the cellular environment, including such conditions as infection with an intracellular pathogen [2], [3], or growth under low oxygen (hypoxic conditions) [4], determines the composition of the exosome. We are still in the early stages of understanding how prokaryotic or eukaryotic intracellular pathogens modulate exosome content and the functions of the exosomes released from the infected cells.

Macrophages are functionally plastic cells with multiple distinct phenotypes regulated by microenvironmental stimuli. The categorization of macrophage function called polarization assigns specific roles to macrophages for such vital processes as an antimicrobial response or tissue repair. While the dichotomy of polarization separates macrophages into classically activated (M1) and alternatively activated (M2) macrophages, the M2 macrophages are further subdivided into M2a, M2b, M2c, and M2d phenotypes that can be distinguished by their expression of separate surface markers and differential production of cytokines with discrete biological functions [as reviewed in [5]]. Prior studies have revealed that the mechanism(s) controlling the initiation of polarization is complex.

However, many of these processes rely on changes in metabolism [6, 7]. A critical pathway that renders M1/M2 polarity relates to the arginine metabolism, where M1 macrophages generally produce iNOS-dependent citrulline and nitric oxide from arginine, in contrast to M2-like macrophages that depend on the arginase metabolic pathway, yielding ornithine and urea as metabolites of the arginine [7]. The critical molecules that influence this polarization in most infections are unknown, although lipopolysaccharides (LPS) and specific Th1 cytokines such as IFN-gamma or TNF promote M1 macrophage activation, while other cytokines such as IL-4 or IL-10 promote M2 macrophage activation or differentiation [8]. Therefore, one could anticipate that the infection from many Gram-negative bacteria is more favorable to initiating M1-like macrophages due to the presence of LPS. However, this polarization might depend on the pathogen andthe type of mouse model used, as studies have shown for *Yersinia enterocolitica* [9] or the pathogen’s growth rate as shown with *Salmonella enterica* serovar Typhimurium [10].

Some studies on the infection of macrophages with the eukaryotic pathogen, *Leishmania*, have explored some of the underlying metabolic contributions that promote macrophage differentiation to M1 or M2 types [11-13]. *Leishmania* parasites are the causative agents of leishmaniasis that can present as cutaneous lesions, diffuse or mucocutaneous lesions, or visceral disease, dependent on the infecting *Leishmania* species. Infection with *L. donovani* causes a visceral disease that presents as splenomegaly, hepatomegaly, and infection of bone marrow cells. Increases in the volume of these organs are suggestive of tissue remodeling characterized in part by an increase in cellularity [14, 15]. As discussed above, macrophages, which are the primary host cells of *Leishmania*, dependent on their activation state, can initiate processes that contribute to tissue remodeling. In addition to the direct effects of the parasite on their host cell, the infection may trigger the release of factors that activate other cells in their vicinity, including bystander macrophages [16, 17]. It is unknown how infected cells communicate with uninfected cells in their vicinity. In our previous studies, we observed that extracellular vesicles (EVs) obtained from *Leishmania donovani*-infected macrophages (LiEV), when incubated with endothelial cells, can induce pathways associated with angiogenesis [18]. We have also performed studies on *Salmonella* infections, where we showed that macrophage responses to infection contribute to the course of infection [19]. Salmonellae are the causative agents of salmonellosis and typhoid fever. Our previous studies described a novel function for EVs in *Salmonella* infection, where EVs from infected macrophages drive M1 polarization in naïve macrophages, leading to the increase in cytokines, such as IL-1 beta or TNF-alpha[20]. Following intranasal delivery of EVs from *Salmonella*-infected cells, they accumulated in the mucosal tissue in the lungs, increasing specific macrophage and DC subpopulations in these areas (2).

To commence comparative studies on the effects of intracellular infections on EV biogenesis and composition, we considered the functions of EVs derived from macrophage infections with either *L. donovani* or *Salmonella*. We show that in the liver of mice infected with *L. donovani*, there is an increase in the macrophage count as the infection progresses. In addition, a protein that had previously been characterized in LiEVs was found to be widely distributed in *L. donovani*-infected livers. This prompted studies to assess the effects of LiEVs on macrophage polarization. In parallel studies, the activation of macrophages by *Salmonella* infection-derived EVs produced by macrophages was evaluated to underscore infection’s effect on EV biogenesis and EV composition.

## 2. Results

### An increase in macrophages in the liver of Leishmania-infected animals

*L. donovani* infection leads to a progressive increase in the weight of the liver and spleen that can be used as a surrogate indicator of the increase in parasitemia within these organs [21, 22]. In the liver, resident macrophages identified by their expression of F4/80 [23] are the primary host cells of *Leishmania* parasites. Within three weeks of infection, granulomas form in the liver, which precedes the eventual drop in parasitemia in the liver [22]. The reduction in parasitemia aside, other outcomes of the infection are the remodeling of the liver characterized by an increase in liver volume and cellularity, which are sustained for many more weeks. Beattie and colleagues [24] observed morphological and other changes in infected and uninfected macrophage populations in infected livers. They then acknowledged that the factors that promoted dynamic changes to the macrophage populations in the infected liver are still unknown. In this study, we sought to gain further insight into the role of the macrophage in this experimental system. Mice were infected with *L. donovani* parasites and monitored over 42 days. Livers were recovered from uninfected mice and mice infected for 20 and 42 days. The livers were weighed, formalin-fixed, and paraffin-embedded. Figure 1a shows a plot of the liver weights over this course of infection. We proceeded to prepare sections from these tissues for analysis. Representative Hematoxylin and Eosin (H&E) stained sections are shown in **Figure 1b**. While there are varying-sized granulomas in the 20-days infected livers, fewer granulomas were seen in the 42-days infected livers, consistent with Murray’s observations[22]. There were no granulomas in the livers of uninfected mice.

**Figure 1.**
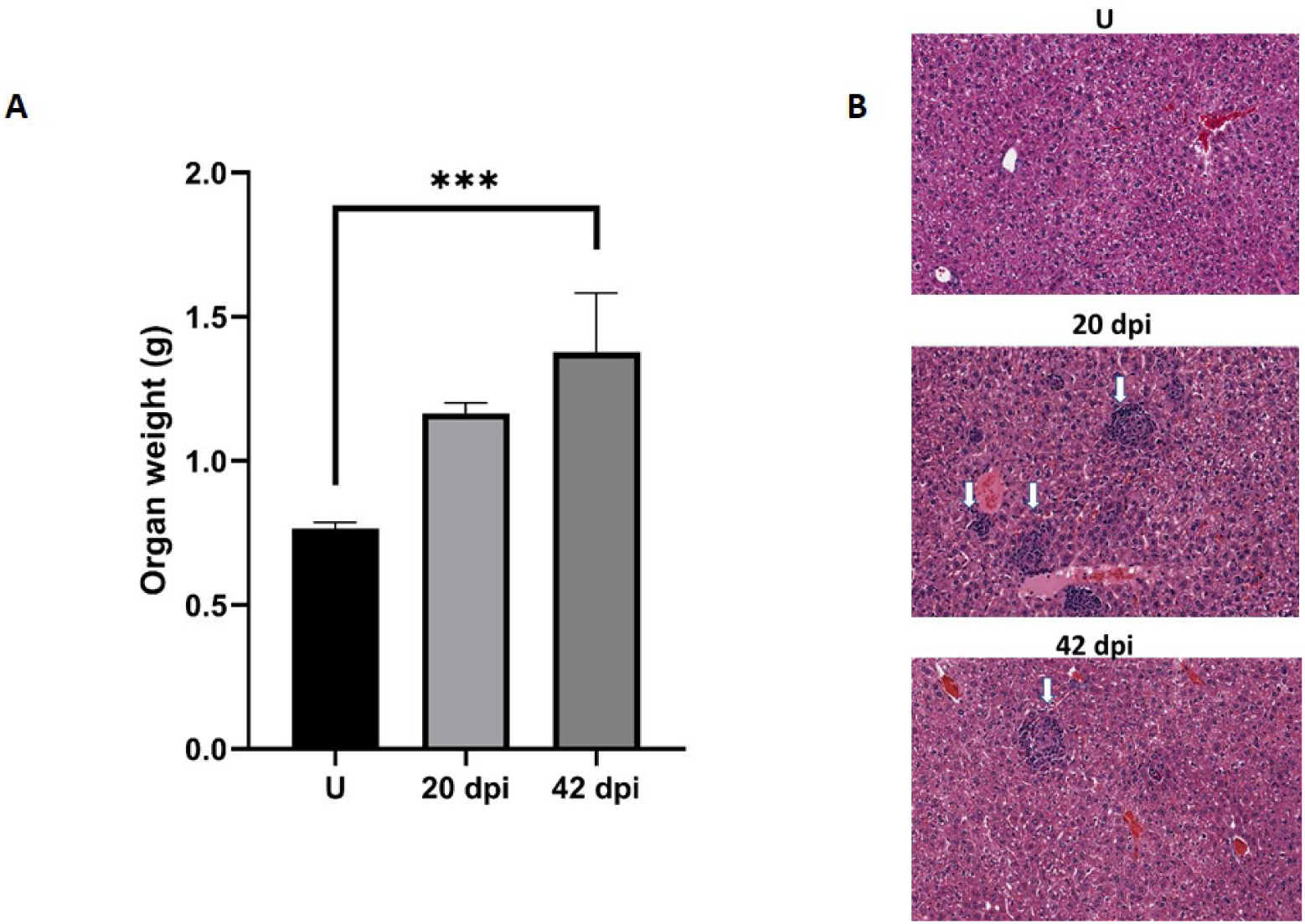
The course of *L. donovani* infection in BALB/c mice. Mice were infected by tail vein injection with metacyclic promastigote forms. Mice were sacrificed after 20 or 42 days. The liver was recovered and weighed before formalin fixation. A) Plot of liver weights. B) Representative H&E sections of liver tissue that was paraffin-embedded and processed for histochemical analyses. White arrows point to granulomas. Each group was composed of 3 animals. ***denotes statistically significant difference p<0.005.

We then evaluated changes to F4/80+ cells in the liver of infected and uninfected mice. Representative images of paraffin-embedded sections prepared from the uninfected liver, 20-day, and 42-day infections and labeled with anti-F4/80 antibodies are shown (**Figure 2A**). There was a sparse distribution of F4/80+ labeled cells in the uninfected liver. In contrast, there was intense F4/80+ labeling of the granulomas and a greater density of F4/80+ labeled cells in livers from 20-days and 42-days infection. Enumeration of F4/80+ cells showed that there was a 3 –4-fold increase in F4/80+ cells at 20- and 42-days post-infection compared to livers from uninfected animals (**Figure 2B**).

**Figure 2.**
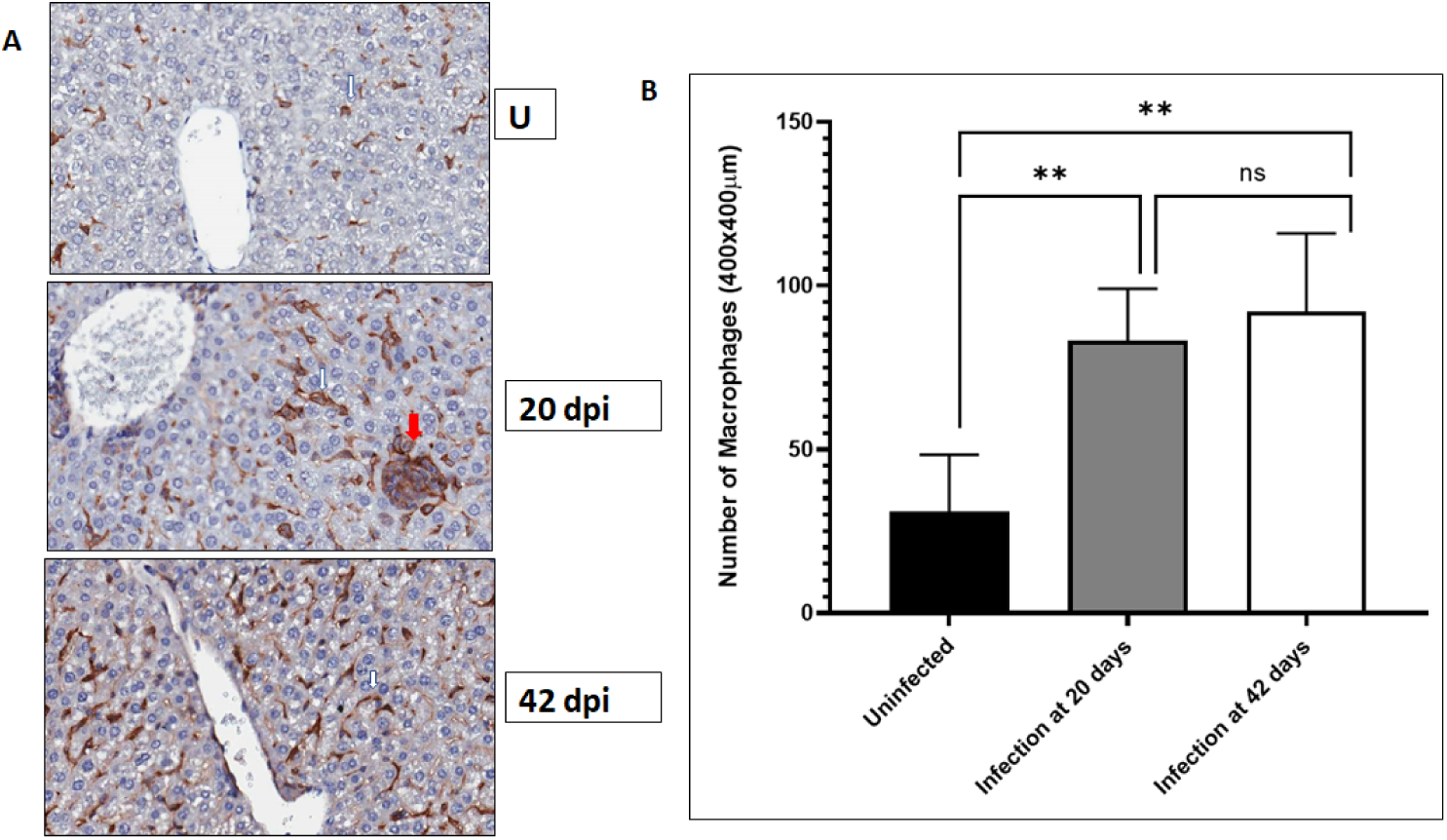
F4/80+ macrophages increase in the liver after *L. donovani* infection. BALB/c mice were infected by tail vein injection of metacyclic *L. donovani* promastigotes. At the indicated times after infection, the livers of infected and uninfected mice were recovered and fixed in formalin and paraffin embedded. Livers sections were prepared and processed for immunohistochemical labeling. F4/80 positive macrophages were labeled with a primary antibody detected with an HRP-conjugated secondary antibody followed by activity of diaminobenzidine (DAB). The sections were counterstained with hematoxylin. A) White arrows point to representative F4/80+ labeled cells; red arrows points to a granuloma. B) F4/80+ cells were enumerated in regions randomly selected through the entire livera nd plotted in GraphPad. Data was compiled from three mice per group. Student t-test was used to establish statistical significance.

**Figure 2.**
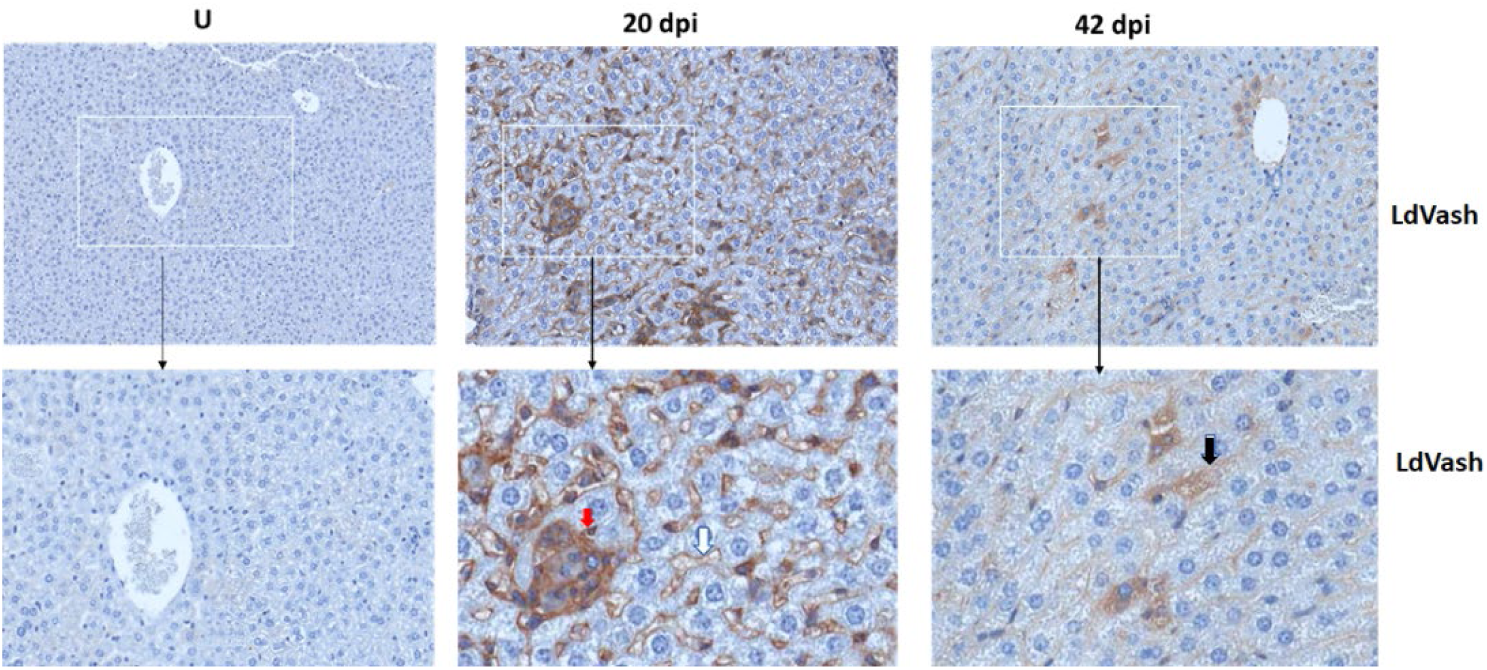
LdVash is released into the liver tissue of infected mice. BALB/c mice were infected by tail vein injection with *L. donovani* parasites. At 20 or 42 days after infection, mice were sacrificed, and their liver was recovered. The tissues were formalin-fixed and then processed for immunohistochemical analyses. Sections from uninfected mice (U) and mice infected for the indicated times were labeled with a custom-made Rabbit anti-LdVash antibody and counterstained with hematoxylin. Regions of interest in the white boxes were magnified. Red arrow arrows point to the label in the granuloma, White arrow points to representative labeling of a sinusoidal capillary. Black arrow points to labeled cell. of an experiment with three mice per group.

### EVs are released from infected cells into the tissue environment

It is likely that infection-derived molecules, including parasite-derived molecules in EVs, play a role in the dynamic changes of macrophages in infected tissues. We had shown that *Leishmania*-infected cells release EVs that contain parasite-derived molecules into the culture medium. To determine whether infected cells in tissues release EVs with parasite-derived molecules, tissue sections from 20- and 42-days post-infection were evaluated for *Leishmania*-encoded Vasohibin (LdVash) distribution. LdVash is a parasite-derived molecule incorporated in EVs released from *Leishmania*-infected cells (LiEVs)[18]. Paraffin-embedded livers from uninfected, 20-days and 42-days old infections were sectioned and labeled with a custom-made peptide antibody to LdVash, as described in the Methods section. Representative images of labeled liver sections are shown (**Figure 3**). In contrast to livers from uninfected mice with no LdVash label, there was a widespread distribution of LdVash in the 20-days post-infection liver. Intense LdVash labeling of granulomas confirmed that infected macrophages, which are the source of LdVash are at the center of granulomas. Interestingly, there was also LdVash labeling of the lining of liver sinusoids, which suggested that EVs and their cargo were widely distributed in the infected liver. In liver sections from 42-days infection, there was evident LdVash labeling of cells, albeit at more reduced levels than the 20-days post-infection livers. It is to be expected that with the reduction in parasite numbers in 42-days infections, there is a reduction in LdVash levels in the liver. Overall, these studies showed that LiEVs and their contents are widely distributed in infected tissues where they can be endocytosed by liver cells, including uninfected bystander macrophages and other liver cells.

**Figure 3.**
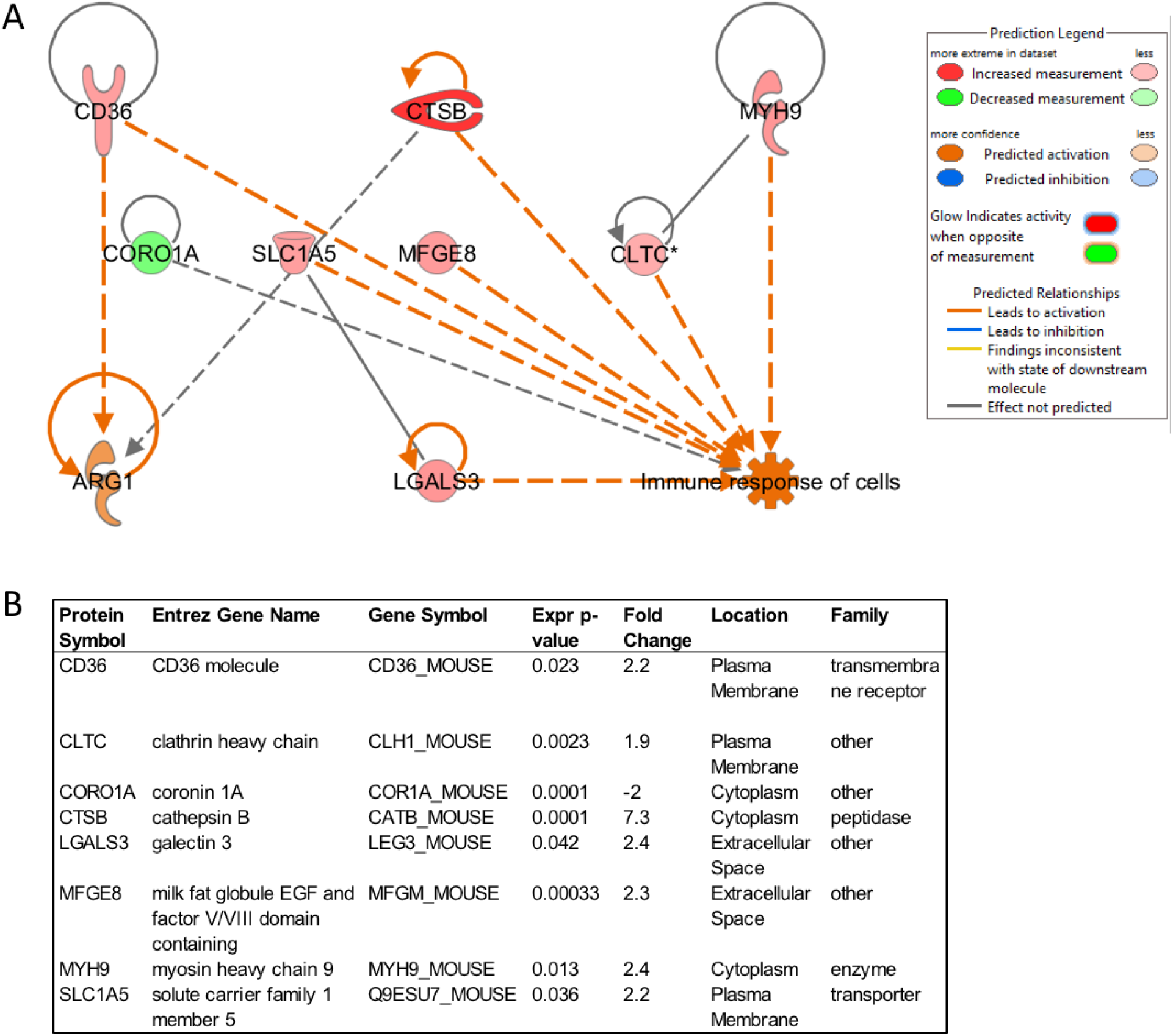
Prediction of downstream molecules and functions regulated by proteins in extracellular vesicles derived from *L. donovani*-infected macrophages (72 hpi). The proteins associated with vesicles produced by uninfected or infected RAW264.7 macrophages were analyzed by quantitative mass spectrometry, followed by analysis using Ingenuity Pathway Analysis. The downstream functions and molecules of vesicular proteins that were upregulated (red) or down-regulated (green) were analyzed, determining the predicted upregulation of Arg1.

### Bioinformatic predictions of the function of molecules in LiEVs

Considering observations that showed that LiEV molecules are widely distributed in infected tissues where macrophages and other cells endocytose these EVs, we sought to predict the potential functions of the EVs based on their composition. We had previously reported on the proteomic composition of LiEVs from RAW264.7 macrophages infected with *L. donovani* parasites[18]. In that study, quantitative mass spectrometry analysis revealed distinct host-derived exosomal proteins with their abundance altered during infection compared to uninfected control. Pathway analysis tools subsequently analyzed the proteins with a protein level differentially regulated by *Leishmania* infection (fold change >2 and p-value <0.05) to identify the mechanisms that could explain the putative function of these EVs. The downstream pathway analysis identified that the upregulation of CD36, CTSB, MYH9, CLTC, MFGE8, SLCS1A5, LGALS3) and downregulation of CORO1A are specific molecules in LiEVs predicted to stimulate the metabolism of cells, represented by the M2 marker ARG1 (**Figure 4**). Arg1 is a metabolic enzyme that catalyzes the hydrolysis of arginine into urea and ornithine [25]. ARG1 is considered a marker of M2 macrophages. In resting macrophages, several stimuli, including cytokines such as IL4 and IL-13, activate STAT6, which in turn upregulates the mRNA and protein levels of ARG1[26]. This bioinformatic analysis led to the testable hypothesis that LiEVs stimulated alternative activation of macrophages. A similar analysis of exosomes derived from *Salmonella*-infected macrophages identified that the EVs generated by this bacterial infection led to increased M1 macrophage polarization[20].

**Figure 4.**
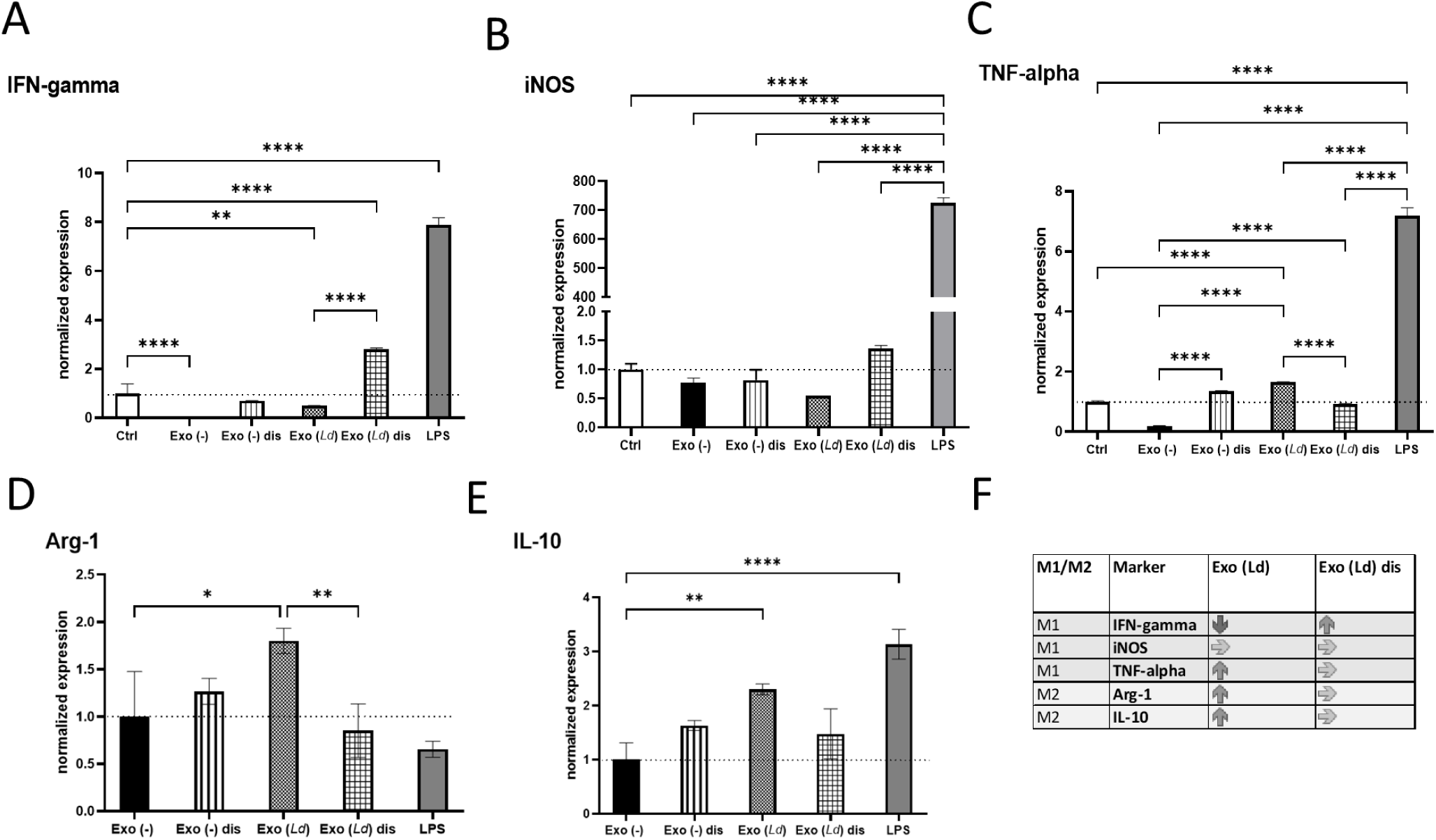
The effect of extracellular vesicles derived from *L. donovani*-infected macrophages on macrophage polarization. RAW264.7 macrophages were infected with *L. donovani* (or not), and the extracellular vesicles were obtained from these cells using the ultracentrifugation method. A portion of the vesicles was disrupted by sonication. Naïve macrophages were then exposed to these vesicles for 24 hours. Media, EVs from uninfected cells, and LPS [100 ng/mL] were used as control treatments. The cells were collected, and RNA purified for qPCR analysis of IFN-gamma (A) iNOS (B), TNF-alpha (C), Arg1 (D), and IL-10 (E) transcripts. One-way ANOVA was used to establish statistical significance. N=6. Exo (-), extracellular vesicles from uninfected cells; Exo (*Ld*), extracellular vesicles from cells infected with *L. donovani;* Exo (-) dis, disrupted extracellular vesicles from uninfected cells; Exo (*Ld*) dis, disrupted extracellular vesicles from cells infected with *L. donovani*; Ctrl, cells not treated.

### Differential activation of primary macrophages by EVs, depending on the pathogen infection

Macrophages exhibit plasticity in their responses to a variety of stimuli. Some stimuli promote macrophages to exhibit gene transcription consistent with classical or M1-type characteristics, while others promote alternatively activated or M2-type characteristics [10, 27]. The specific stimuli in each infection promoting such macrophage polarization are still being studied. As discussed above, the bioinformatic analysis of the LiEVs proteome had led to the prediction that these EVs would likely promote alternative activation of macrophages, based on the predicted increase in the Arg-1 protein (**Figure 3)**. Hence, we aimed to determine the effect of EVs on macrophage activation. LiEVs from RAW264.7 macrophage infected with *L. donovani* parasites were incubated with peritoneal exudate cells prepared from BALB/c mice. Macrophage expression of prototypic M1-type and M2-type activation markers was determined after 24-hrs. LiEVs and EVs from uninfected cells did not induce the expression of IFN-gamma, iNOS, or TNF-alpha, which contrasted with LPS that induced conspicuous expression of these markers (**Figure 4**). Instead, LiEVs and not EVs from uninfected cells induced a statistically significant increase in the M2 markers, Arg-1, IL-10 compared to the treatment with exosomes from uninfected cells. Disruption of LiEVs before incubation with macrophages mitigated their capacity to activate the induction of Arg-1 and IL-10. Although LPS did not activate Arg-1 expression, we observed induction of IL-10 expression in response to LPS, which is consistent with observations by others [28]

To further address the critical role of the composition of EVs on the macrophage responses that are elicited, EVs prepared from RAW264.7 macrophages infected with *S*. Typhimurium were evaluated alongside LiEVs. As expected, EVs derived from *S*. Typhimurium-infected cells induced significant iNOS and TNF-alpha levels, comparable to levels of these markers induced by LPS treatment of cells. EVs from *S*. Typhimurium-infected cells did not increase Arg-1 or IL-4R transcripts. This is consistent with observations on macrophage activation in bystander cells or in cells harboring non-growing *Salmonella* [10] In contrast, LiEVs did not induce iNOS or TNF alpha transcription but increased Arg-1 and IL-4R (**Figure 5 A-D**). As another finding, the vesicles derived from *Salmonella*-infected macrophages did not cause increased expression of iNOS when the vesicles were disrupted by sonication, but the effect on TNF-alpha transcript expression was less pronounced (**Figure 5 E-F**). Measurement of soluble cytokine levels confirms this observation (Supplemental Figure 1). It is worth noting that differences in responses elicited by these EVs underscore the effects of their composition on macrophage activation.

**Figure 5.**
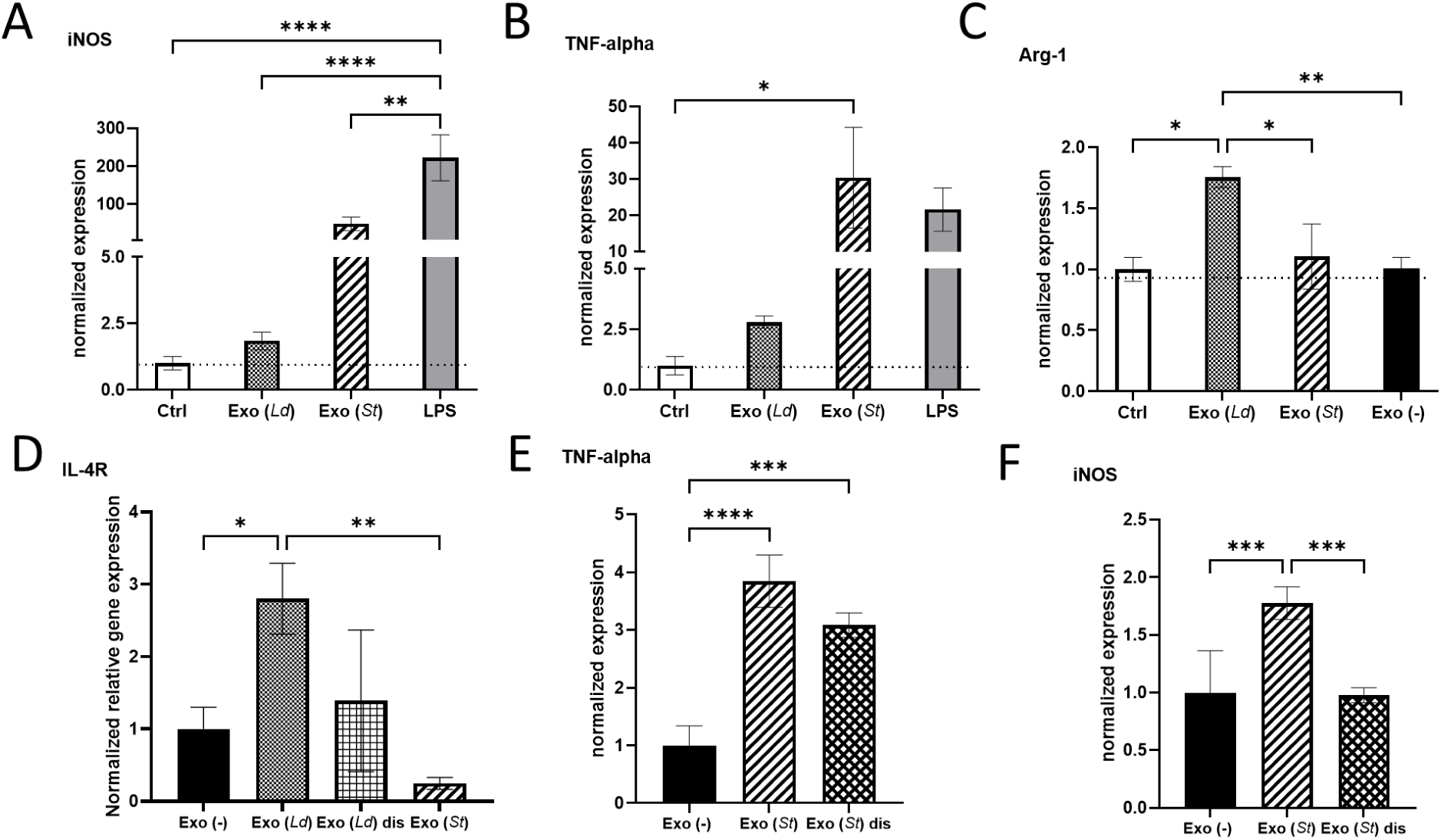
The effect of extracellular vesicles derived from *L. donovani*-infected macrophages on macrophage polarization. RAW 264.7 macrophages were infected with *L. donovani* (72 hpi) or *Salmonella enterica* ser. Typhimurium (MOI 5:1, 24 hpi). The EVs obtained from these cells were used to treat primary macrophages for 24 hours. Media (Ctrl) or EVs from uninfected cells were used as negative controls, and LPS treatment 100 ng/mL was used as a control treatment. The cells were collected and RNA purified for qPCR analysis of iNOS, TNFa, Arg1, and IL-4R. One-way ANOVA for multiple comparison tests was used to establish statistical significance. N=6. Exo (-), extracellular vesicles from uninfected cells; Exo (*Ld*), extracellular vesicles from cells infected with *L. donovani;* Exo (*St*), extracellular vesicles from cells infected with *S*. Typhimurium; Exo (*St*) dis, disrupted extracellular vesicles from cells infected with *S*. Typhimurium, Ctrl, cells not treated.

### M1 and M2 macrophages in *Leishmania*-infected liver

Studies described above showed that LdVash, a prototypic LiEV molecule, is conspicuously distributed in infected mouse livers, especially at 20 days after infection. As was described earlier, LdVash is not only detected within infected F4/80+ cells but also in bystander F4/80+ cells. Next, F4/80+ cells in the liver were examined in the extent to which they displayed characteristics of classically activated or alternatively activated cells. Towards this goal, the expression of M1 or M2 signature molecules was ascertained in these cells. Tissue sections from 20-days infected livers were evaluated for expression of iNOS (M1 signature) or IL-4Rα (CD124) (M2 signature. Although iNOS labeling was sparse, it was primarily detected in the center of granulomas (**Figure 6A**). In contrast, IL-4Rα labeled F4/80+ cells were more widely distributed in the infected livers.

**Figure 6.**
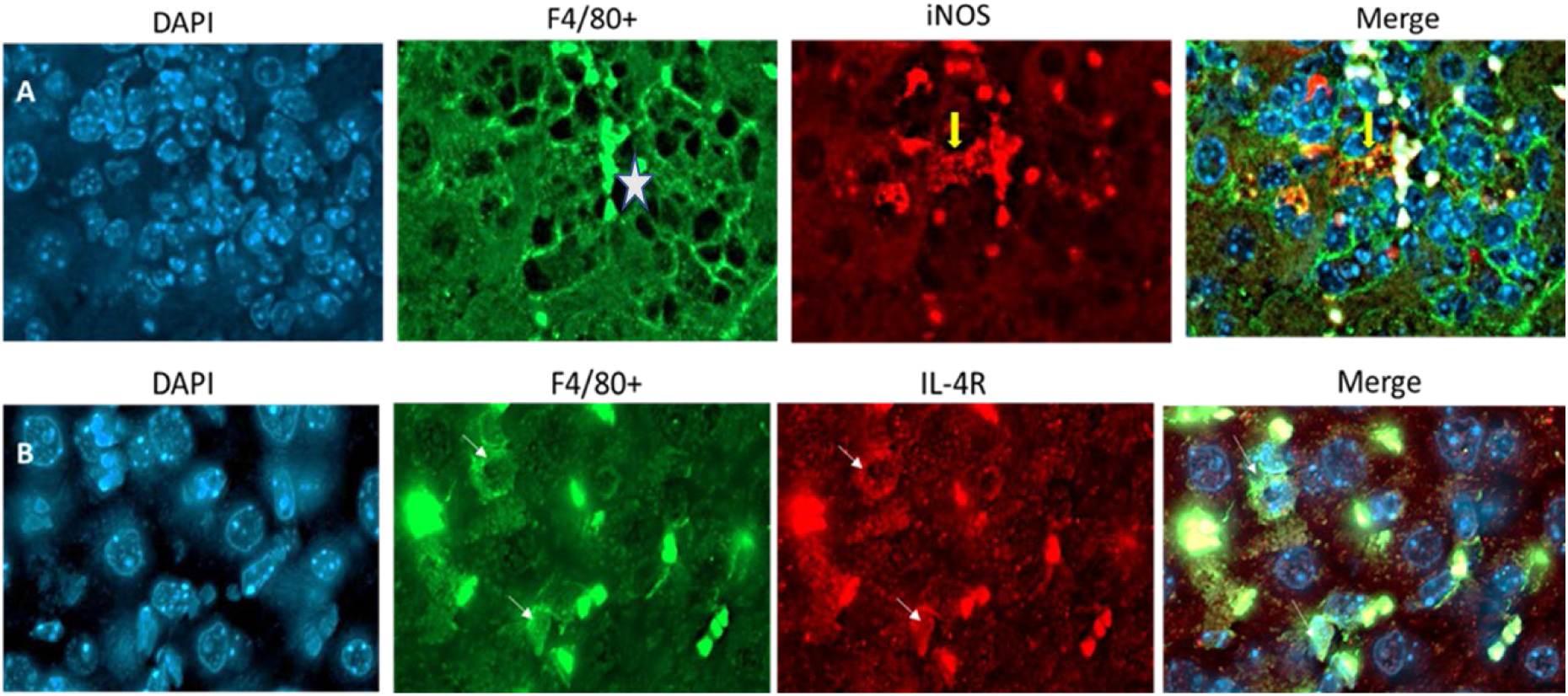
Macrophages within infected tissues display M1 marker restricted to granuloma and more widespread expression of M2 marker. BALB/c mice were infected by tail vein injection with *L. donovani* parasites. At 20 days after infection, mice were sacrificed, and their liver recovered. The tissues were formalin-fixed and then processed for immunohistochemical analyses. Sections from the infected were labeled with anti-F4/80 and either Inos (A) or IL-4R (B). Yellow arrow in A, points to specific labeling of iNOS within a granuloma (white star is in the center of the granuloma). White arrows in B point to specific Il-4R labeling. Images are representative of 3 mice in infected group.

## Discussion

In infections of *Leishmania donovani*, the liver, spleen, and bone marrow are parasitized. Several studies have shown that parasitemia in the liver eventually drops in experimental infections after achieving peak levels. This disease progress contrasts with the situation in the spleen, where parasite numbers and tissue volume continuously increase[29] Granuloma formation in the liver precedes and foretells the drop in parasite numbers even though liver volume and cellularity do not change significantly. Kupffer cells, the liver’s resident macrophages, are the primary hosts of *Leishmania* parasites in the liver [30]. Kupffer cells in the liver are characterized by the expression of F4/80+[23]. The current understanding of the contributions of Kupffer cells to the progress of the infection in the liver is incomplete. Both infected and uninfected macrophages may play complementary roles in the infection. Our studies found a 3-to-4-fold increase in F4/80+ macrophages in the livers of infected mice over a 42-days infection course (**Figure 2**). The infected F4/80+ cells, which were found to be widely distributed in the livers of 20-day (3 weeks) infection, were at the center of granulomas, which is consistent with the observations by others [21, 22]. Although the number of granulomas was significantly reduced by 42 - days post-infection, macrophage numbers remained high. Several studies have reported on the cytokine milieu in *Leishmania*-infected liver and spleen tissues, and some experiments have shown the role of cytokines in the remodeling of the infected tissues[31-33]. However, it is not known what role infection-derived molecules, including parasite molecules, may play in promoting cell activation in infected tissues.

To follow up on our previous studies that characterized EVs derived from *Leishmania*-infected macrophages (LiEVs) and identified several parasites molecules in these EVs [18], we sought to determine whether LdVash, a prototypic parasite molecule in LiEVs, was released into infected livers. Immunolabeling with a custom-made anti-peptide antibody to LdVash revealed that it was widely distributed in the liver in 20-day old infections. In addition to labeling granulomas that contain infected macrophages at their center, LdVash was also seen in other cells, including cells lined livers sinusoids (Figure 3). This result suggested that infected F4/80+ Kupffer cells are potentially modulated by parasites and that infection-derived products could functionally modulate uninfected bystander macrophages and other non-immune cells that ingest LiEVs. Our previous studies showed that LiEVs could activate endothelial cells in surrogate angiogenesis assays, including tube formation and the release of angiogenesis-promoting factors [18]. Macrophages are critical cells of the innate immune system, serving as primary phagocytic cells that produce ROS for pathogen destruction [34], cells that present antigens on MHC I and MHC II receptors to T-cells [35], and releasing cytokines that signal to other cells to infiltrate into the infected region [36]. However, macrophages are plastic cells that can modify their metabolism and hence the functional properties, depending on the stimulants they encounter [8]. As such, macrophages can affect the immediate environment by producing cytokines, leading to the induction of proinflammatory responses in the case of M1 macrophages or enhanced phagocytosis and tissue healing properties in the case of M2 macrophages [36]. Signals such as LPS, IFN-λ, TNF-α, or IL-1β induce differentiation into M1 macrophages, displaying increased proinflammatory gene transcription and bactericidal properties[37]. For example, infection with *S*. Typhimurium leads to an LPS-dependent M1 macrophage profile in murine macrophages [38]. Such M1 polarized macrophages release IFN-λ, TNF-α, or IL-1β and upregulate inducible nitric oxide synthase (iNOS) in naïve cells to promote bacterial killing [reviewed in [39]].

Polarization to M2, or alternatively activated macrophages, is mediated by IL-4 [40] and generally occurs after M1 responses to decrease inflammation [41] and maintain homeostasis [42]. IL-4 induces M2 type gene transcription such as Arg-1, which mediates cell growth via metabolizing L-arginine into urea and L-ornithine, which are crucial for generating proline and polyamines necessary for collagen production represses ROS generation by downregulating iNOS expression[43, 44]. IL-10 is typically secreted from M2 macrophages and acts as an anti-inflammatory cytokine [45, 46], restricting tissue damage during infection [47]. The balance between M1 and M2 macrophages appears critical for proper immune responses by balancing proinflammatory and anti-inflammatory responses[36, 41, 42]. It is unknown whether the vesicles produced by macrophages during Leishmania infection lead to M2 polarization, but they certainly stimulate the gene expression consistent with M2 polarization. In contrast, the vesicles produced during *Salmonella* infection stimulate the transcription of M1 polarization markers [2]. Since the vesicles can migrate to tissues far from the infection site, there is a likelihood that the vesicles produced during infection can stimulate macrophages far from the site of their generation.

## 3. Materials and Methods

### Bioinformatic analysis

Ingenuity Pathway Analysis software (Qiagen) was used for protein network analysis of exosomal proteins, focusing on the analysis of proteins with different abundance upon infection with *L. donovani* at 72 hpi compared with exosomes isolated from uninfected RAW 264.7 macrophages. Activation of specific downstream proteins and functions was identified and measured by Z-score higher than 2/-2, where relevant canonical pathways were overlayed.

### Cell culture

Murine macrophage cell line RAW264.7 (ATCC# TIB-71, ATCC, USA) was cultured in DMEM media that was supplemented with 10% fetal bovine serum (FBS) and 100 μg/ml Penicillin/Streptomycin (Life Technologies Inc., USA) at 37°C and 5% CO_2_. Peritoneal exudate cells (PECs) t were obtained 4 days after intraperitoneal injection of thioglycolate into BALB/c mice.

*L. donovani* wild type (MHOM/S.D./62/1S-CL2_D_) was obtained from Dr. Nakhasi’s lab (FDA) and cultivated in M199 media (Sigma M0393) containing 15% FBS, 0.1 mM Adenosine, 0.1 mg/mL folic acid, 2 mM glutamine, 25 mM HEPES, 100 units/mL penicillin/100 μg/mL streptomycin (Gibco 15140122), 1X BME vitamins (Sigma B6891), and 1 mg/mL sodium bicarbonate with pH 6.8 at 26°C

*Salmonella enterica* serovar Typhimurium strain UK-1 χ3761 (wild-type) was cultured at 37°C in lysogeny broth (LB) media and shaken at 200 rpm (rotations per minute). After the overnight culture, the bacterial cultures were diluted in new LB media to reach the optical density at 600 nm (OD_600_) of 0.05. This culture was grown until OD_600_ reached 0.50, the mid-logarithmic phase. The bacteria were then washed in 1 mL of phosphate-buffered saline (PBS) and centrifuged at 6,000 x g for 10 minutes at 37°C before bacterial cultures were used for infections.

### Infection conditions

RAW264.7 cells were washed with pre-warmed PBS, and incomplete growth media containing no FBS or antibiotics were added for 60 minutes before infection with *S*. Typhimurium at a multiplicity of infection (MOI) of 5:1. The overnight culture of *S*. Typhimurium was diluted as described above and cultivated at 37°C until it reached the mid-logarithmic phase. The bacteria were then added onto the cells for 2 hours, after which time the culture media were removed, cells were washed with PBS, and media containing gentamicin (100 μg/mL) were added onto the cells for 1 hour. Finally, the media containing lower gentamicin (20 μg/mL) and exosome-free heat-inactivated FBS were added to the cells for the remaining time of infection. Based on our previous studies, the total infection time for *S*. Typhimurium was 24 and 48-hours, based on our previous studies [2, 19].

For *L. donovani* infections, RAW264.7 macrophages were plated on 100 mm culture dish containing sterile glass coverslips at a concentration of 5 × 10^6^ cells per dish, and cells adhered overnight at 37°C with 5% CO_2_ before infection with metacyclic promastigotes. To enrich for metacyclic parasites, peanut agglutination (PNA) was performed on stationary-phase wild-type promastigote cultures[48]. Briefly, 4-d-old cultures of *L. donovani* parasites were washed twice and resuspended in incomplete DMEM at a 2 × 108 parasites/ml concentration. PNA was then added to the parasites at a final concentration of 50 μg/ml and incubated at room temperature for 15 min. The parasites were then centrifuged at 200*g* for 5 min to pellet agglutinated parasites. The supernatant was then collected, and the PNA-metacyclic parasites were washed twice, resuspended in complete DMEM, and counted for infection. Parasites were then added to macrophage dishes at a ratio of 20:1 (parasites: macrophage).

### Extracellular vesicle purification

As described previously, the extracellular vesicles were isolated from the cell culture media by an ultracentrifugation method [2, 18]. Cell culture supernatant was briefly collected and filtered through a 0.22-micron polyethersulfone (PES) filter. PBS containing 1 mM PMSF and 1X protease cocktail protease inhibitor cocktail (EDTA-free; Roche, USA) was added. The filtrates were centrifuged at the following conditions: 10 min at 500 × g, 10 min at 2,000 × g, and 40 min at 16,000 × g, and after each step, they were moved to a new tube. Finally, the supernatant was ultracentrifuged for 180 min at 100,000 × g using SW 32 Ti rotor and Optima XPN ultracentrifuge (Beckman, USA). The pellets were washed with PBS containing the protease inhibitor cocktail, and additional centrifugation was carried out at 100,000 × g to wash the pellet. The EVs were resuspended in a sterile PBS containing protease inhibitor cocktail (Roche, USA). Next, the vesicles were counted using Nanoparticle Tracking Analysis (NTA, NanoSight LM10) to measure the vesicle concentration and the hydrodynamic diameter.

### Treatments of cells with EVs

Peritoneal exudate cells (PECs) were treated with EVs or controls for 2, 24, or 72 hours. EVs were obtained from uninfected RAW 264.7 macrophages, *Salmonella* infected RAW 264.7 macrophages, *Leishmainia donovani* infected RAW 264.7 macrophages, or *Leishmania amazonensis* infected RAW 264.7 macrophages at 5*10^9^ or 1*10^9^ EV particles per mL of media. Control treatments included exosome depleted DMEM media and LPS 100 ng/mL. Following treatment, supernatants and cell pellets were collected and stored at -80°C. Cell pellets were lysed

### qPCR analysis

The transcription of murine iNOS, Arg-1, IL-4R, IFN-gamma, and TNF-alpha was analyzed using the primers described previously [2], while transcripts of IL-4R and IFN_gamma were performed using the primers described in **Supplemental Table 1**. RNA was extracted from macrophages using a Qiagen RNeasy Mini Plus extraction kit. The cDNA was generated from the isolated RNA using Maxima First Strand cDNA synthesis kit (Thermo Fisher), and the expression of genes was measured by using a two-step quantitative real-time polymerase chain reaction (RT-qPCR) using MXP3005 instrument (Bio-Rad) and SYBRGreen reagents (Bio-Rad). Primer sequences were validated by melt-curve analysis.

### Mouse infections and Histochemistry

Six to eight-week-old groups of female BALB/c mice (3 per group) were inoculated intravenously (tail vein) with 1 × 10^7^ stationary phase *L. donovani*. The liver of each mouse was recovered and weighed on days 20 and 42 post-infection. Fresh liver tissues were placed in cassettes, fixed in 10% formalin, and embedded in paraffin (FFPE). Histochemistry was carried out by the Molecular Pathology Core at the University of Florida. For hematoxylin and Eosin staining, tissue sections (4 μm) were deparaffinized with xylene, and the tissue sections were rehydrated in a graded series of ethanol solutions. The rehydrated tissues were stained with hematoxylin (Richard-Allan Scientific, 7212) for 2 minutes, incubated with clarifier 2 (Richard-Allan Scientific, 7402) for 30 seconds, followed by incubating with a bluing reagent (Richard-Allan Scientific, 7301) for 1 minute, then incubated one minute in 80% ethanol before staining with eosin (Richard-Allan Scientific, 71311) for 1 minute. In between the application of each reagent, the slides were washed with running water. Finally, the H&E stained slides were dehydrated in a graded ethanol series, dipped in xylene, and the cover slipped. For immunohistochemistry. 4 mm sections were deparaffinized and rehydrated by serially passing through xylene and graded ethanol changes. For single-stained slides, sections were subjected to heat-induced antigen retrieval in 10mM Citra pH 6 and blocked with avidin, biotin, and goat serum. They were incubated overnight at 4°C with primary rabbit antibodies against F4/80 or LdVash. After washing, tissues were labeled with Mach2 Gt x Rabbit HRP polymer (Biocare Medical, Walnut Creek, CA), the DAB chromogen (Biocare Medical, Walnut Creek, CA), and CAT hematoxylin counterstain (Biocare Medical, Walnut Creek, CA). Whole slides were scanned using an Aperio CS2 Scanscope (Leica/Aperio, Vista, CA). For double-stained slides, FFPE liver sections on slides were heated in EDTA buffer (10 mM Tris, 1 mM EDTA solution, pH 9.0 - Thermo Fisher) for antigen retrieval, followed by 3% H_2_O_2_. After washing, tissues were blocked with Background Sniper (BioCare Medical), followed by incubating with rabbit anti-mouse F4/80 overnight. After washing, the slides were incubated for 1hr at RT with secondary horse anti-rabbit FITC IgG. After washing, the slides were subsequently incubated with either anti-iNOS or anti-CD124 (ThermoFisher) overnight. Slides were washed and incubated with the appropriate secondary antibody. Slides were washed, then counterstained with DAPI, and viewed using an immunofluorescence microscope (Zeiss, Axioskop 2 mot plus, Oberkochen, Germany). Data were analyzed by Image J (National Institutes of Health).

## 4. Acknowledgments

Histochemistry was performed by the UF Molecular Pathology Core. This work was supported by U. S. Public Health Grant R03 AI-135610 (MJE), R01 AI158749-02 (MJE), R56 - AI143293 (PEK).

## 5. Figure legends

**Supplemental Table 1.**
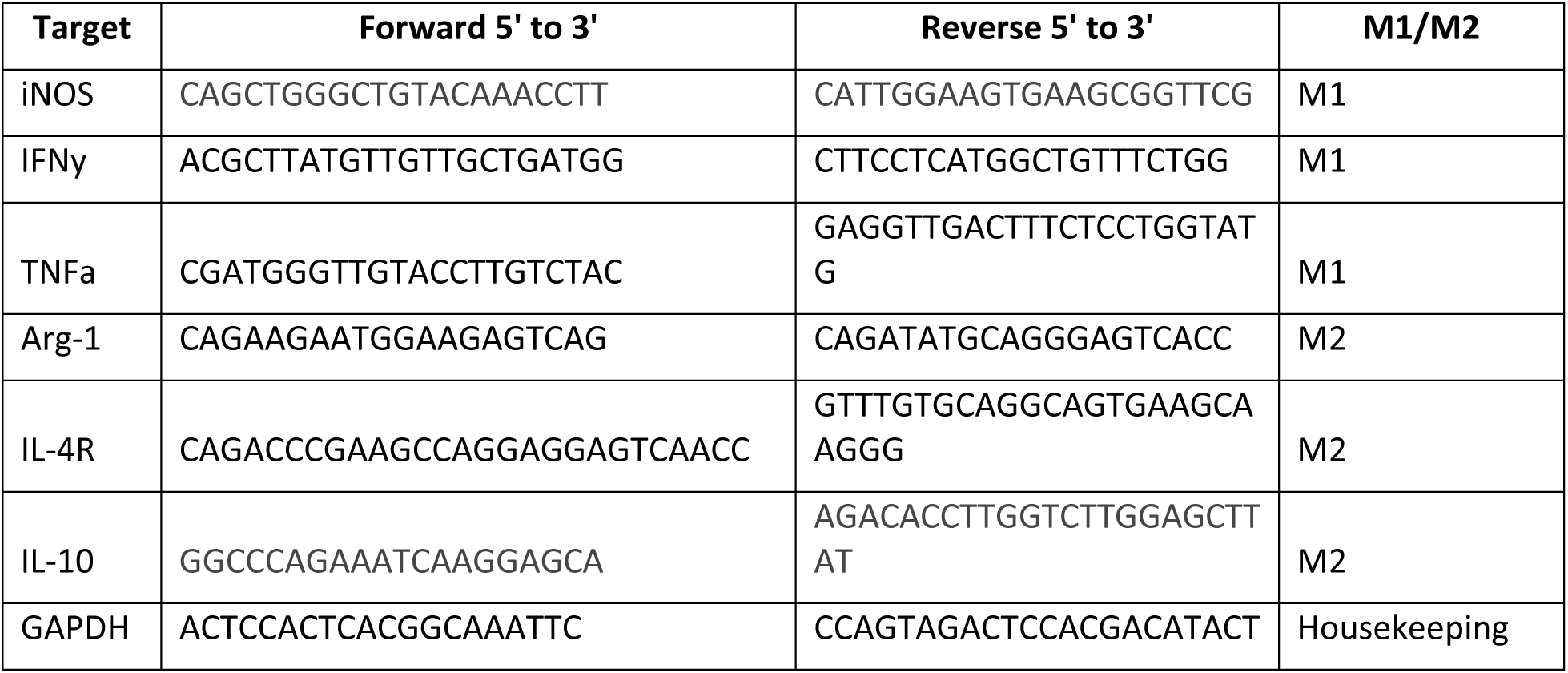
Primers.

## Supplementary figures

**Figure S1.**
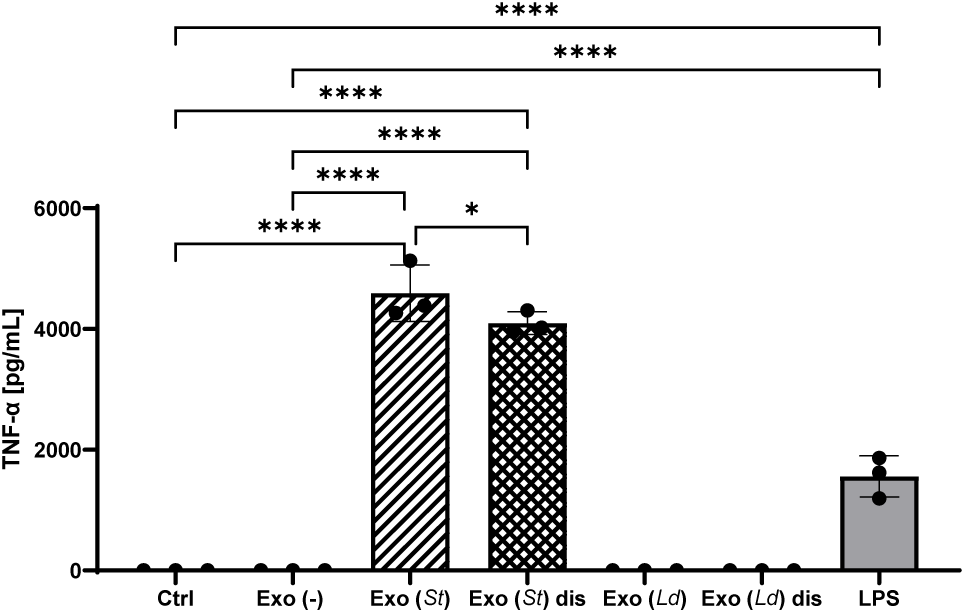
The effect of extracellular vesicles derived from *L. donovani*- or *Salmonella* Typhimurium-infected macrophages on TNF-alpha release. RAW 264.7 macrophages were infected with *L. donovani* (72 hpi) or *Salmonella enterica* ser. Typhimurium (MOI 5:1, 24 hpi). The EVs obtained from these cells were used to treat primary macrophages for 24 hours. Media (Ctrl) or EVs from uninfected cells were used as negative controls, and LPS treatment 100 ng/mL was used as a control treatment. The cell culture supernatants were collected and processed for anti-TNF ELISA. One-way ANOVA for multiple comparison tests was used to establish statistical significance. N=3. Exo (-), extracellular vesicles from uninfected cells; Exo (*Ld*), extracellular vesicles from cells infected with *L. donovani;* Exo (*St*), extracellular vesicles from cells infected with *S*. Typhimurium; Exo (*St*) dis, disrupted extracellular vesicles from cells infected with *S*. Typhimurium, Ctrl, cells not treated.

